# No evidence for cognitive impairment in an experimental rat model of knee osteoarthritis and associated chronic pain

**DOI:** 10.1101/2022.08.22.504732

**Authors:** Sara Gonçalves, Gareth J. Hathway, Stephen G. Woodhams, Victoria Chapman, Tobias Bast

## Abstract

Although chronic pain states have been associated with impaired cognitive functions, including memory and cognitive flexibility, the cognitive effects of osteoarthritis (OA) pain remain to be clarified. The aim of this study was to measure cognitive function in the mono-iodoacetate (MIA) rat model of chronic OA-like knee pain.

We used young adult male Lister hooded rats, which are well suited for cognitive testing. Rats received either a unilateral knee injection of MIA (3mg/50µL) or saline as control. Joint pain at rest was assessed for up to 12 weeks using weight-bearing asymmetry, and referred pain at a distal site using determination of hindpaw withdrawal thresholds (PWT). The watermaze delayed-matching-to-place (DMP) test of rapid place learning, novel object recognition (NOR) memory assay and an operant response-shift and -reversal task were used to measure memory and behavioural flexibility. Open field locomotor activity, startle response, and prepulse inhibition (PPI) were also measured for comparison.

MIA-injected rats showed markedly reduced weight-bearing on the injured limb, as well as pronounced cartilage damage and synovitis, but interestingly no changes in PWT. Rearing was reduced, but otherwise locomotor activity was normal and no changes in startle and PPI were detected. MIA-injected rats had intact watermaze DMP performance, suggesting no substantial change in hippocampal function, and there were no changes in NOR memory or performance on the operant task of behavioural flexibility. Our finding that OA-like pain does not alter hippocampal function, unlike other chronic pain conditions, is consistent with human neuroimaging findings.

## 1. INTRODUCTION

Chronic pain can impair cognitive function, in a way that appears to depend on the chronic pain condition, possibly reflecting differences in pain intensity or different pathophysiological mechanisms [1,40,52]. It has been suggested that chronic pain reduces cognitive resources by diverting them to pain-related stimuli or by impairing structure and function of key brain regions, including the hippocampus and prefrontal cortex (PFC) [41,52]. Although additional factors, including pain medication and co-morbidities, are also likely to contribute to cognitive impairment associated with chronic pain in patients [40,52], experimental studies have shown impairments in cognitive functions associated with hippocampus and PFC in rodent models of neuropathic [31,33,41] and inflammatory pain [37,45,46].

Osteoarthritis (OA) is a very common chronic pain condition, estimated to affect 250 million people worldwide, with knee OA having the highest prevalence [21]. Recent longitudinal studies show that OA pain is associated with cognitive impairments, reflected in measures of verbal recall and fluency, subtraction ability, and attention [24,27]. Moreover, grey matter changes [4] and functional reorganisation [5,23] across cortical and subcortical regions have been identified in OA patients. However, unlike chronic back pain and complex regional pain syndrome, OA patients do not have a significant loss of hippocampal volume [41]. Experimental models of OA have been widely used to study the mechanisms of OA pain behaviour (see refs in [18]). Although one study reported impaired novel object recognition (NOR) memory in the murine mono-iodoacetate (MIA) model of OA pain [43], the impact of rodent models of OA pain on distinct cognitive functions has yet to be addressed. Such rodent models make it possible to study the impact of chronic pain separately from confounding factors, including comorbidities or pain medication.

The aim of this study was to measure a range of cognitive functions, which have been linked to the hippocampus and PFC, in the MIA rat model of OA-like knee pain. We used Lister hooded rats as these are more suitable for cognitive testing than the albino rat strains commonly used in pain research [20,29,66]. To assess distinct cognitive functions, study 1 used the watermaze delayed-matching-to-place (DMP) task of rapid place learning, which is highly sensitive to hippocampal dysfunction [7,10,61], and study 2 used the NOR memory test, which depends on medial temporal lobe structures, including hippocampus and perirhinal cortex [12,65], and an operant task of behavioural flexibility, which depends on PFC function [9]. Cognitive testing was complemented by measurements of pain behaviour (weight-bearing asymmetry and von Frey test of paw-withdrawal threshold (PWT)) and of sensorimotor processes (acoustic startle response and its prepulse inhibition [PPI] and open-field locomotor activity).

## 2. METHODS

### 2.1. Rats

We ran two studies, using two separate cohorts of young adult male Lister hooded rats (250-275g, about 8 weeks old upon arrival, Charles River, UK): study 1 (watermaze DMP test of rapid place learning) used a total of 16 rats, and study 2 (NOR memory test and operant test of behavioural flexibility) used 32 rats (Figure 1). We only included male rats, because the parameters of the MIA model and of the behavioural assays were chosen based on previous studies in males. Also, in our hands, female rats have shown less robust performance on the watermaze DMP task than males (Charlotte Taylor and Tobias Bast, unpublished observations). Nevertheless, we recognise that our focus on male rats is an important limitation.

**Figure 1.**
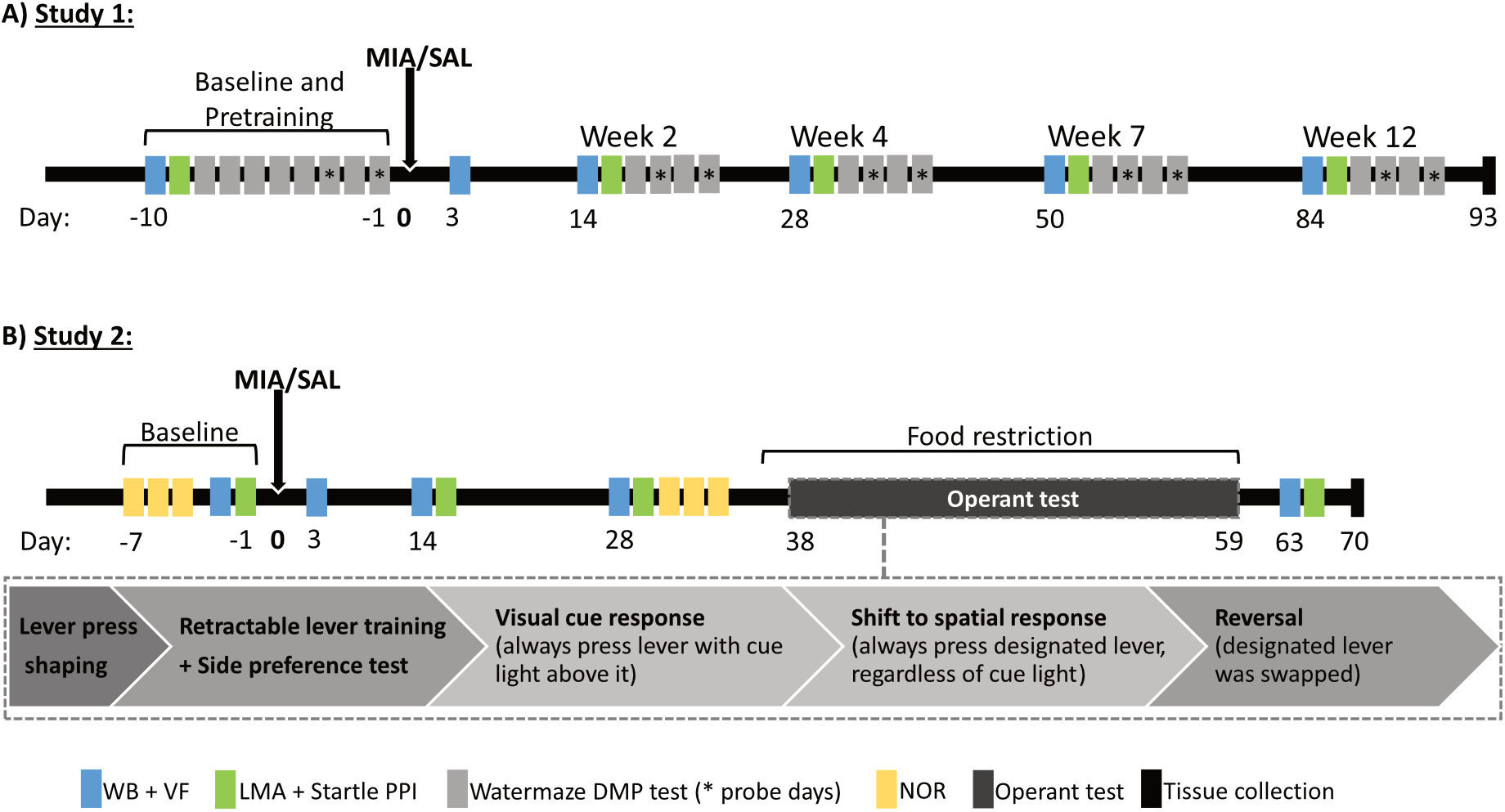
Overview of study 1 and study 2. **A)** Timeline for study 1 – watermaze DMP test of rapid place learning: Before MIA model induction, baseline pain behaviour and sensorimotor measures were collected and pre-training and baseline measurements on the watermaze DMP task were completed. Aer model induction (day 0), rats were tested on the watermaze DMP task in weeks 2, 4, 7 and 12 to longitudinally evaluate the impact of OA-like chronic knee pain on hippocampus-dependent rapid place learning performance. Pain behaviour (weight-bearing and Von-Frey tests) and sensorimotor processes (locomotor acvity and startle and its PPI) were measured in the same weeks, before watermaze DMP testing. Rats were euthanised on day 93, and knees were collected for post mortem analysis. **B)** Timeline for study 2 – NOR memory test and operant test of behavioural flexibility: Before MIA model induction, baseline measures of NOR, pain behaviour and sensorimotor processes were collected. Aer model induction (day 0), NOR memory was re-assessed on day 29-31 after model induction. Aer this, rats were put on food restriction from day 35 to 59 and rats underwent testing on the food-reinforced operant task of behavioural flexibility from day 38 to 59. Pain behaviour and sensorimotor processes were measured at several me points following model induction, before and after NOR and operant testing. Rats were euthanised on day 70, and knees were collected for post mortem analysis. DMP, delayed-matching-to-place; LMA, locomotor acvity; MIA, mono-iodoacetate; NOR, novel object recognition; PPI, prepulse inhibition; VF, Von-Frey; WB, weight-bearing.

Rats were group housed, 4 per cage in individually ventilated cages (“double decker” cages -462 × 403 × 404 mm; Tecniplast, UK), in temperature- and humidity-controlled conditions (21 ± 1.5°C, 50 ± 8%) and with an alternating 12/12 h light/dark cycle (lights on at 7 A.M). Rats had food and water available *ad libitum*, except when testing on the food-reinforced operant task in study 2. Following arrival in the animal unit, rats had a period of acclimatisation (3-7 days), during which they were habituated to handling by the experimenter for a few days, before any experimental procedures started. All behavioural testing was carried out during the light phase. Studies had the approval of the University of Nottingham’s Animal Welfare and Ethical Review Board (AWERB) and were conducted in accordance with the requirements of the UK Home Office Animals Scientific Procedures Act (1986) and the International Association for the Study of Pain. Procedures are reported in compliance with the ARRIVE guidelines [47].

### 2.2. Induction of the monosodium iodoacetate (MIA) model of osteoarthritis pain

Rats received a single intra-articular injection of either 3 mg/50 µL MIA (Sigma, UK) or 50 µL 0.9% saline (control group) through the infra-patellar ligament of the left knee, using a 30-gauge 8-mm hypodermic needle, under brief isoflurane anaesthesia (isoflurane 2.5 – 3% in 1L/min O2) [32]. Health and welfare checks were performed immediately after anaesthetic recovery, daily for 5 days, and weekly thereafter. The MIA dose was selected based on previous studies conducted in albino rats (see refs in [18]) and a pilot study in young adult male Lister hooded rats. In our pilot study, young adult Lister hooded rats injected into the knee with a dose of 1 mg/50 µL MIA did not show robust pain behaviour and had only weak OA-like knee pathology, whereas 3 mg/50 µL of MIA produced marked pain behaviour and knee pathology (pilot data not shown). Rats were allocated randomly to the treatment groups, with half of the rats in each cage receiving MIA treatment and the other half saline. The experimenter was blinded to the treatment allocations throughout data collection and analysis.

### 2.3. Pain behaviour and sensorimotor testing

In both study 1 and 2, baseline pain and sensorimotor behaviours were assessed prior to model induction and then at several subsequent time points (Figure 1). Pain behaviour was assessed using two behavioural tests: joint pain at rest was assessed by weight-bearing asymmetry using an incapacitance tester (Linton Instrumentation, Diss, UK) [8]; and referred pain at a site distal to the injured joint was assessed via determination of mechanical PWT using the up/down method for both ipsilateral and contralateral hind paws [11,14]. Weight-bearing asymmetry was calculated as [ipsilateral g/(ipsilateral g + contralateral g)]. As the intervals between successive von Frey hairs are logarithmic, PWTs were reported as log(von Frey hair values in g) [38].

Locomotor activity and the acoustic startle response and its PPI are basic sensorimotor processes, modulated by forebrain regions, including hippocampus and PFC [6]. Altered performance in these tests may point to changes in these forebrain regions and would also need to be considered when interpreting any changes in performance on the cognitive tests (because gross sensorimotor effects may confound cognitive testing). Locomotor activity was assessed in open-field chambers (39.5cm x 23.5cm x 24.5cm) with two 4×8 photobeam configurations, one close to the bottom of the cage and the other one close to the top of the cage (Photobeam Activity System; San Diego Instruments, US), as previously described [49]. Two consecutive breaks of adjacent bottom photobeams generated a horizontal locomotor count, and breaks of photobeams of the top configuration generated rearing counts. Open field sessions lasted 30 min. Immediately after locomotor testing, acoustic startle response and PPI of the startle responses were measured, using the SR LAB Startle Response System (San Diego Instruments, US), as described previously [49]. Test sessions lasted a total of 23 min, throughout which rats were exposed to a 62-dB (A) background noise. After 5-min acclimatization, rats were exposed to a series of 10 startle pulses (40-ms, 120-dB (A) broad-band bursts) presented alone, so that the startle response could habituate to a relatively stable level of startle reactivity for the remainder of the session. In the second block, 50 trials were presented to measure PPI. There were 5 different trial types, each one presented 10 times in pseudo-random order and with a variable inter-trial interval (10-20s): pulse-alone trials and four types of prepulse-plus-pulse trials, in which a weak 20-ms prepulse [72, 76, 80, or 84 dB(A)] preceded the startle pulse by 100 ms. The percentage of PPI (%PPI) induced at each prepulse intensity was calculated as: %PPI = [(mean amplitude on pulse-alone trials – mean startle amplitude on prepulse-plus-pulse trials) / (mean startle amplitude on pulse-alone trial)] X 100.

### 2.4. Measurements of learning and memory and of behavioural flexibility

Detailed descriptions of the apparatus and procedures can be found in the Supplementary Material.

#### 2.4.1. Watermaze DMP task of rapid place learning

The watermaze delayed-matching-to-place (DMP) task is highly sensitive to hippocampal dysfunction [7,35,48,61], requiring rats to learn within one trial the daily changing place of a hidden platform to escape efficiently from a circular pool of water, which is surrounded by prominent visual cues for spatial orientation. Rats perform four daily trials. During trial 1, rats rapidly learn the novel location of the hidden platform, and then on subsequent trials use the place memory to efficiently locate the hidden platform. Trial 2 is occasionally run as “probe trial”, during which the platform is removed to monitor the rats’ search preference for the zone containing the platform (correct zone). In study 1, rats were pre-trained on the watermaze DMP task, and their baseline performance was measured before MIA model induction. After model induction, watermaze DMP performance was evaluated at weeks 2, 4, 7 and 12 (Figure 1A). Methods were adapted from our previous studies [35]. Search preference for the platform area and its vicinity during probes was used as the main performance measure of rapid, 1-trial, place memory performance. Percentage of time searching the correct zone was calculated as: [time in correct zone (s)/total time in all 8 zones (s)] x 100%. By chance, i.e., if the rats were searching randomly, this value would be 100%/ 8 = 12.5%. Additionally, path lengths to reach the platform perimeter were recorded for all trials, with steep reduction from trial 1 to trial 2 indicating acquisition of 1-trial place memory. Swim speed was also measured during the 60-s probe trials, to examine potential motor impairments in the MIA-injected rats. For analysis, search preference and swim speed was averaged across the two probes at each testing time point (on day 6 and 8 for baseline testing and the two probes conducted as part of the 4-day testing blocks at each post-operative testing time point) and path length measures for trial 1 to 4 were averaged across all four days at each testing time point (days 4 to 8 of pretraining for baseline values and all four days of the 4-day test blocks at all post-operative testing time points).

#### 2.4.2. NOR memory test

The NOR memory test exploits rats’ innate tendency, without use of training or reinforcement, to explore novel stimuli, including novel objects, more than familiar stimuli [17,50]. During the test, rats are presented with a novel object and a familiar object, i.e. an object that they had the opportunity to explore during a preceding sample phase. Rats will typically spend more time exploring the novel object, reflecting recognition memory of the familiar object explored during the sample phase. Brain areas required for simple NOR memory include medial temporal lobe regions, especially the perirhinal cortex [65], and the hippocampus, particularly if the retention delay between sample phase and test is greater than 10 min [12]. In study 2, NOR memory was assessed before MIA model induction, and on day 29-31 after model induction (Figure 1B), using a procedure with a 24-h retention delay between sample phase and test, which was adapted from previous work [50]. Time exploring each object (direct contact with or active exploration of the object by directing the nose towards the object from a distance of less than 1 cm) was measured in both the sample phase (two identical objects) and the test phase (one familiar and one novel object). The discrimination ratio was calculated as follows: D = total time exploring novel object/ (total time exploring novel object + total time exploring familiar object).

#### 2.4.3. Operant test of behavioural flexibility

Behavioural flexibility is the ability to switch stimulus-response patterns rapidly, as reflected by response shifts or reversals [62], and can be assessed in rats using a food-reinforced operant test [9]. Shifts refer to a subject beginning to make responses to new or previously irrelevant stimuli (e.g., from pressing the lever indicated by a cue light to pressing either the left or right lever in an operant box), whereas reversals refer to changing a response from a previously rewarded to a previously non-rewarded stimulus within the same category (e.g., from left to right lever). Previous studies suggest that behavioural flexibility, as measured on the operant task, depends on the PFC and subcortical regions, including ventral striatum [9]. In study 2, following MIA model induction (from day 38 to 59), rats were trained first to acquire a lever press response to a cue light in order to receive food reward, before being tested for their ability to shift this response to a spatial response (pressing either the left or right lever to receive food) and then to reverse this spatial response (start to press the opposite lever to receive food reward) (Figure 1B). Operant testing procedures were adapted from previous studies [9]. The analysis of this test focused on trials to criterion, the percentage of correct responses, and percentage of omissions.

### 2.5. Assessment of OA-like knee pathology

At the end of study 1 (day 93 after model induction) and 2 (day 70 after model induction), rats were anesthetised with an overdose of sodium pentobarbitone (Dolethal, Vetoquinol; 2mL intraperitoneal), and then transcardially perfused with 0.9% saline followed by 4% paraformaldehyde in 0.9% saline.

Both knees were collected, decalcified, stained and histologically scored to confirm development of OA-like pathology via established methods [25,34,53]. Briefly, knee joints were stored in the fixative for 72h after collection, transferred to an ethylenediaminetetraacetic acid + 7.5% polyvinylpyroolidene solution for decalcification (6-7 weeks), and then split in the frontal plane [28]. Trimmed joints were mounted in paraffin wax and 5µm sections cut using a microtome. Haematoxylin and Eosin [60] and Safranin-O-Fast Green [60] histological stains were used to confirm OA-like damage to the knee joints. Cartilage damage was scored on a scale of 1-5 for severity with a multiplier of 1-3 for area coverage, to give a composite score out of 15. Osteophytes were scored 1-3 based on size (1 = mild <40µm; 2 = moderate, 40-160µm, 3 = severe, >160µm) [25,34,53].

### 2.6. Sample sizes and statistical analysis

The target sample size for both studies was 32 (n=16 per group), so group differences corresponding to an effect size of Cohen’s d=1 could be detected with a power of about 80%, using an independent t-test (2-tailed, p<0.05). In study 1, we used a sequential design [44]. The first series of testing with half the target sample size revealed that there were no substantial group differences in the main memory measures, and indicated that completion of the second series to achieve the target sample size would not reveal significant group differences. Therefore, study 1 was terminated after completion of the first series [44]. Moreover, one MIA rat was excluded from study 1 due to a physiological abnormality (swelling in the non-injected right hind paw, possibly due to fighting), preventing completion of behavioural data collection. Therefore, in study 1, data analysis was based on n=8 saline and n=7 MIA rats. In study 2, 32 rats were tested in parallel, and no rats had to be excluded from data analysis, resulting in n=16 rats in both the saline and MIA group.

GraphPad Prism 8 and IBM SPSS Statistics 24 were used to prepare the graphs and perform the statistical analysis. All data were normality tested using D’Agostino-Pearson test prior to subsequent analyses. The different measurements were analysed, using analysis of variance (ANOVA) with group (MIA or saline) as between-subjects factor and testing day as repeated measures/within-subjects factor (2-way ANOVA) and with task block (open field and startle data), prepulse intensity (PPI data), trial (watermaze and operant data) and object (novel vs. familiar, NOR data) as within-subjects factors, as appropriate (3-way ANOVA). Baseline measures were not included in the ANOVA analysis, but were analysed separately, using independent t-tests or 2-way ANOVA (with prospective group as between-subjects factor and task block, prepulse intensity or trial as within-subjects factor), to ensure that there was no significant group difference at this stage. Trials-to-criterion measures from the operant task and knee pathology measures were compared between groups, using independent t-test or the Mann-Whitney U test, if the normality assumption was not met. P<0.05 was accepted as significance threshold for all analyses.

## 3. RESULTS

### 3.1. Pain behaviour and sensorimotor measurements: MIA knee injections induced weight-bearing asymmetry and decreased rearing activity

MIA-treated rats placed markedly less weight on the injured leg compared with saline-injected rats across all testing time points in both study 1 (Figure 2A) and 2 (Figure 2D), resulting in significant weight-bearing asymmetry (study 1 – group: F_(1,13)_=66.22, p<0.0001; time: F_(4,52)_=3.32, p<0.0001; time x group: F_(4,52)_<1; study 2 – group: F_(1,30)_=33.89, p<0.0001; time: F_(3,90)_=0.71, p=0.55; group x time: F_(3,90)_=0.21, p=0.89). However, there were no differences in PWT between MIA-, and saline-injected rats across all testing time points following injection in both study 1 (Figure 2B) and study 2 (Figure 2E) (study 1 – group: F_(1,13)_=1.0, p=0.33; time: F_(4,52)_=1.66, p=0.18; time x group: F_(4,52)_<1; study 2 – group: F_(1,30)_=1.68, p=0.20; time: F_(2.9,87.4)_=1.62, p=0.19; time x group: F_(3,90)_<1).

**Figure 2.**
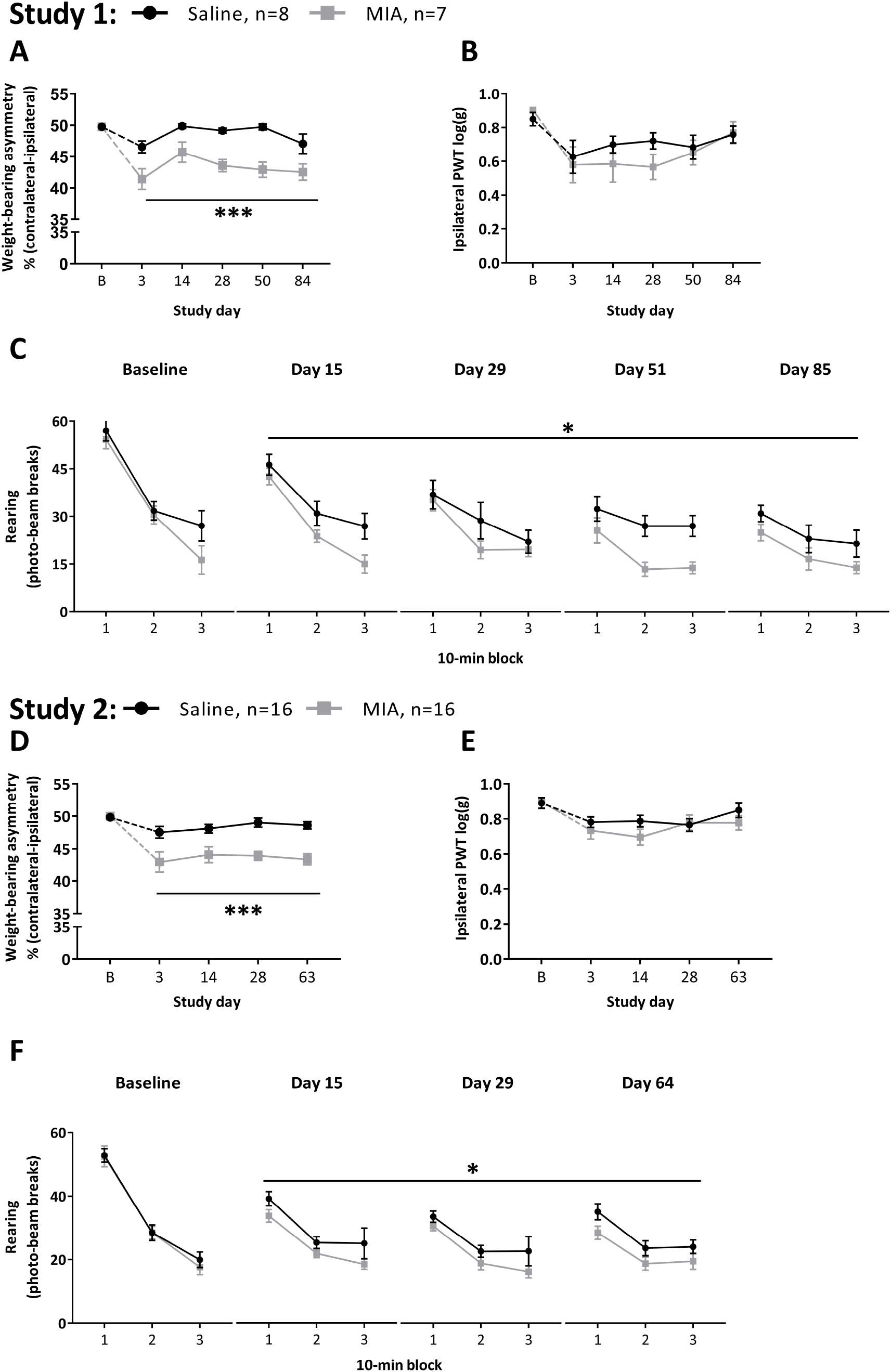
MIA-induced knee pain in Lister hooded rats: weight bearing asymmetry and reduced rearing, but no significant change in paw withdrawal threshold (study 1 and 2). Rats were injected with either 3mg/50µl MIA (▪, grey squares) or 50µl saline (•, black dots) into the le knee. In both studies, MIA injection induced marked weight bearing asymmetry across all testing me points after model induction (study 1, **A**; study 2, **D)**, whereas no changes were observed in the von-Frey test of paw withdrawal threshold (study 1, **B;** study 2, **E**). Rearing was significantly reduced in MIA-injected rats compared with saline controls across the complete 30-min open field test sessions and across all testing me points after model induction (study 1, **C;** study 2, **F**). Data are presented as mean±SEM. * and *** indicate significant ANOVA main effect of group across all testing me points after model induction (2-way ANOVA, p < 0.05 and p < 0.0001, respectively). B, baseline; MIA, mono-iodoacetate; PWT, paw withdrawal threshold.

MIA-treated rats did not show altered horizontal locomotor activity in study 1 (group: F_(1,13)_<1, all interactions involving group: F<1), but there was a trend for reduced locomotor activity in study 2 (group: F_(1,30)_=3.65, p=0.07, all interactions involving group: F<1.05, p>0.05) (Figure S1). MIA-treated rats showed significantly less rearing than saline controls in both study 1 (Figure 2C) and study 2 (Figure 2D) (study 1 – main effect of group: F_(1,13)_=4.86, p=0.046; all interactions involving group: F<1.9, p>0.18; study 2 – main effect of group: F_(1,30)_=5.08, p=0.03; all interactions involving group: F<1). MIA-treated rats did not exhibit altered startle reactivity or PPI (all main effects or interactions involving group: F

<2.14, p>0.15, except for a trend towards a main effect of group on startle response in study 2, F_(1,30)_=2.91; p=0.10, reflecting a numerical decrease in the startle response in rats injected with MIA, which was already present at baseline, i.e. before knee injections, and persisted across all testing time points) (Figures S2 and S3).

### 3.2. MIA-induced OA-like knee pain pathology

MIA-treated rats showed marked OA-like knee pathology (Figure S4). MIA-treated rats had lateral and medial tibial plateau loss of cartilage integrity (study 1-t_(13)_=4.63, p=0.0005; study 2-U=24, p<0.0001), as well as increased synovial inflammation (study 1-U=0, p=0.0003; study 2-U=48, p=0.0001) compared to saline-injected rats. There was a trend for MIA-injected rats to show a higher number of osteophytes in study 1 (U=14, p=0.08) and a significant increase in study 2 (t_(29)_=2.51, p=0.02).

### 3.3. Intact hippocampus-dependent rapid place learning performance after MIA model induction

At baseline testing, both prospective groups showed similar marked pathlength reductions from trial 1 (encoding) to trial 2 (retrieval) (main effect of trial: F_(3,90)_=12.5; p<0.0001; interaction prospective group X trial: F_(3,90)_<1), as well as similar (t_(30)_<1) and markedly above-chance (t_(15)_>3.68, p<0.002) search preference for the correct zone when trial 2 was run as a probe (Figure 3). After MIA treatment, both groups showed virtually identical performance patterns in terms of path length across all testing time points (main effect of group and interaction involving group: all F<1), with marked path length reductions from trial 1 to 2 (trial: F_(3,84)_=80.19; p<0.0001) (Figure 3A). Both groups also showed similar search preference for the correct zone when trial 2 was run as a probe trial (main effect and interaction involving group: F<1.8; p>0.20), and search preference was highly above chance in both groups at all testing time points (saline: t_15_>2.69, p<0.018; MIA: t_15_>3.17, p<0.006) (Figure 3B). The only significant effect of group on performance measures was a significant interaction of group and time point in the ANOVA of swim speed data (F_(3,84)_=8.87; p<0.0001), reflecting that MIA rats had slower swim speed than control rats at the last two testing time points (Figure 3C). More specifically, control rats slightly increased their swim speed from week 7, whereas the swim speed of MIA rats remained stable.

**Figure 3.**
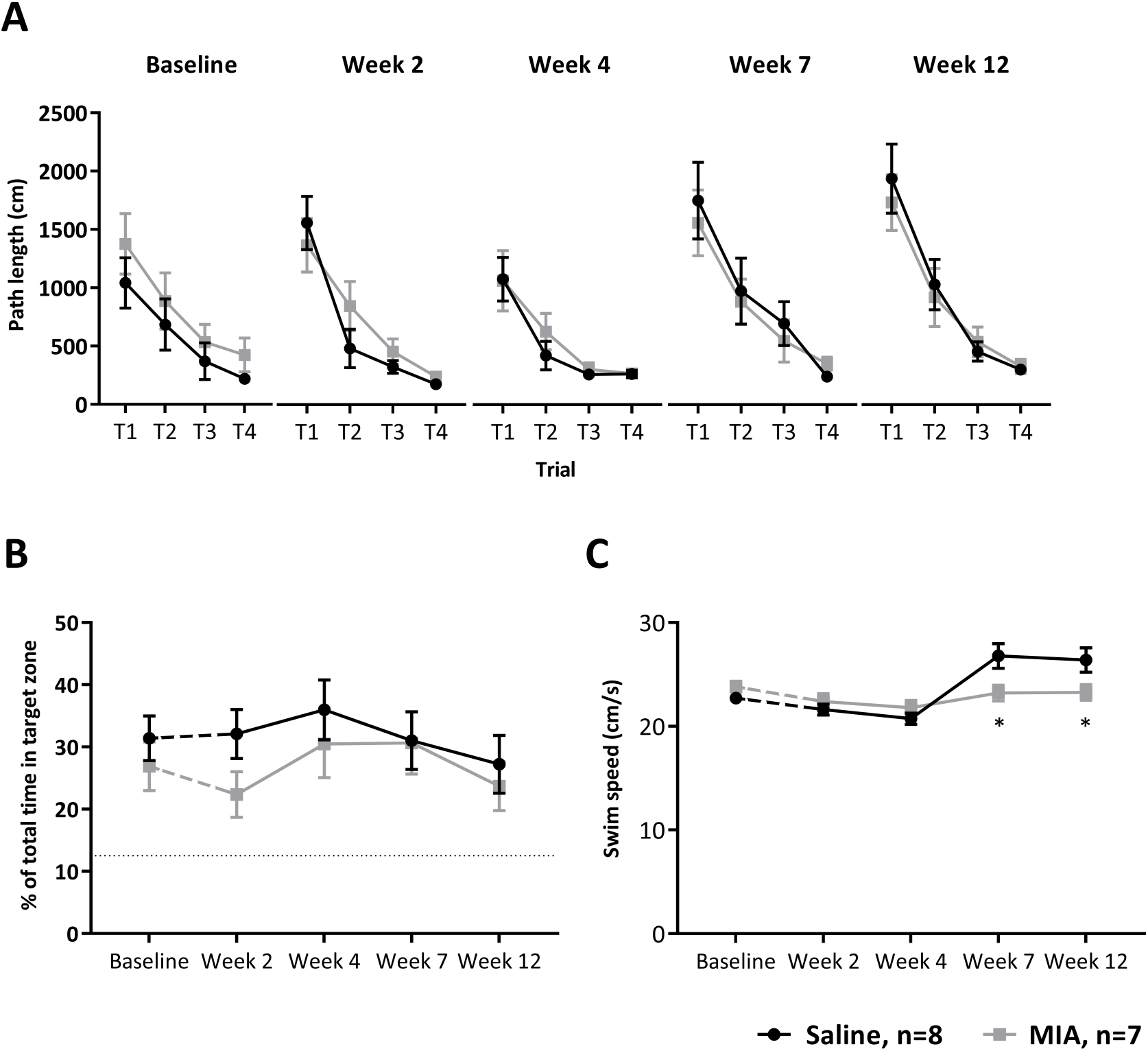
Intact rapid place learning performance on the watermaze Delayed-Matching-to-Place (DMP) task following MIA model induction. Rats were injected with either 50ul of 3mg of MIA (▪, grey squares) or 50µl saline (•, black dots) in the le knee. They were tested on the watermaze DMP task at baseline, i.e. before the knee injections, and then during weeks 2, 4, 7 and 12 after the knee injections. **A**) Path lengths to reach the platform across the four daily trials (T1 to T4) to a novel location show marked reduction across trials, including from trial 1 to 2, reflecting rapid place learning, which was not affected by MIA injection. **B**) Search preference for the target zone when trial 2 to the daily new goal location was run as probe, with the escape platform unavailable, was markedly above chance (12.5%, stippled line) across all testing me points, refleticting 1-trial place learning; search preference was unaffected by MIA injection. **C**) Swim speed was similar in both groups during baseline and in week 2 and 4 following injection, but was increased in saline compared to MIA-injected rats in week 7 and 12 following knee injection. Data are presented as mean±SEM. *p<0.05 MIA vs. saline, following group X interaction in 2-way ANOVA.

### 3.4. Intact NOR memory after MIA model induction

At baseline and after model induction, exploration time was similar in both groups (F_(1,30)_<1), and both groups showed a similar preference for the novel object (main effect of object: F_(1,30)_>27, p<0.0001; no group X object interaction: F<1) (Figure 4A). The discrimination ratio was not altered in the MIA-treated group (group: F_(1,30)_=2.27, p=0.14), and was significantly different from chance in both groups at baseline and after model induction (t>2.41, p<0.03) (Figure 4B).

**Figure 4.**
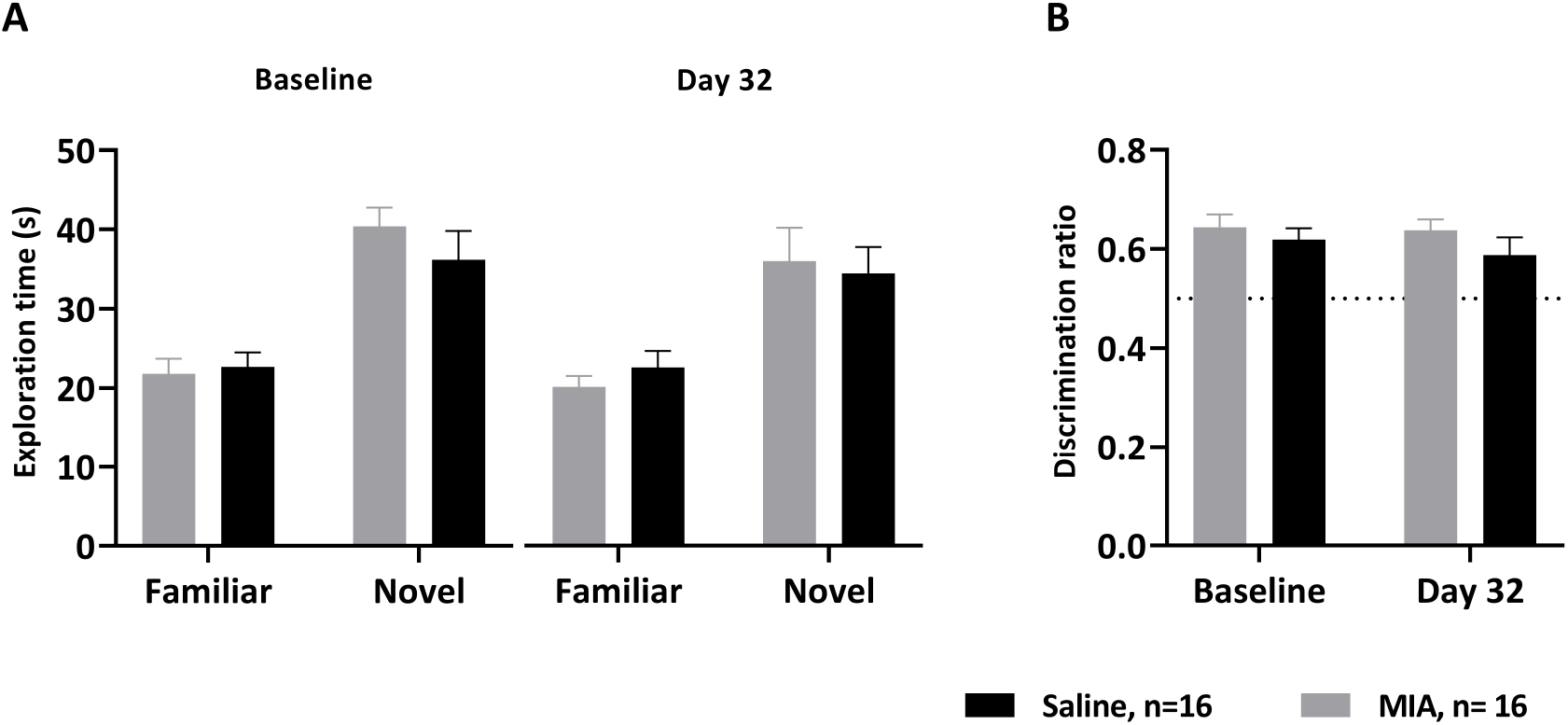
Intact novel object recognition (NOR) memory following MIA model induction. Rats were injected with either 50ul of 3mg of MIA (grey) or saline (black) in the le knee. They were tested on the NOR memory task at baseline, i.e. before the knee injections, and 32 days after the knee injections. Both groups showed similarly marked recognition memory for the familiar object at both testing me points. This was reflected by increased **A**) exploration me for the novel, compared to the familiar, object and **B**) an above-chance (0.5, stippled line) discrimination ratio. Data are presented as mean±SEM.

### 3.5. Intact performance on the operant task of behavioural flexibility after MIA model induction

Both MIA-treated and control rats completed the pretraining on the operant task efficiently. MIA treatment did not impair any aspect of the operant testing. Both groups learned the cue discrimination similarly well, showing similar trials to reach the criterion (t_(30)_<1; Figure 5A), % of correct responses (t(30)=1.35, p=0.2; Figure 5B) and % of omissions (median, percentiles: saline, 0.5, 0-1.75; MIA, 0, 0-1; U=109.5, p=0.46, data not shown). At the visual discrimination stage, 4 rats (3 saline, 1 MIA) failed to achieve the criterion of 10 successive correct responses within three 150-trial sessions (highlighted by the ellipses in Figure 5A) and, therefore, were excluded from further testing.

**Figure 5.**
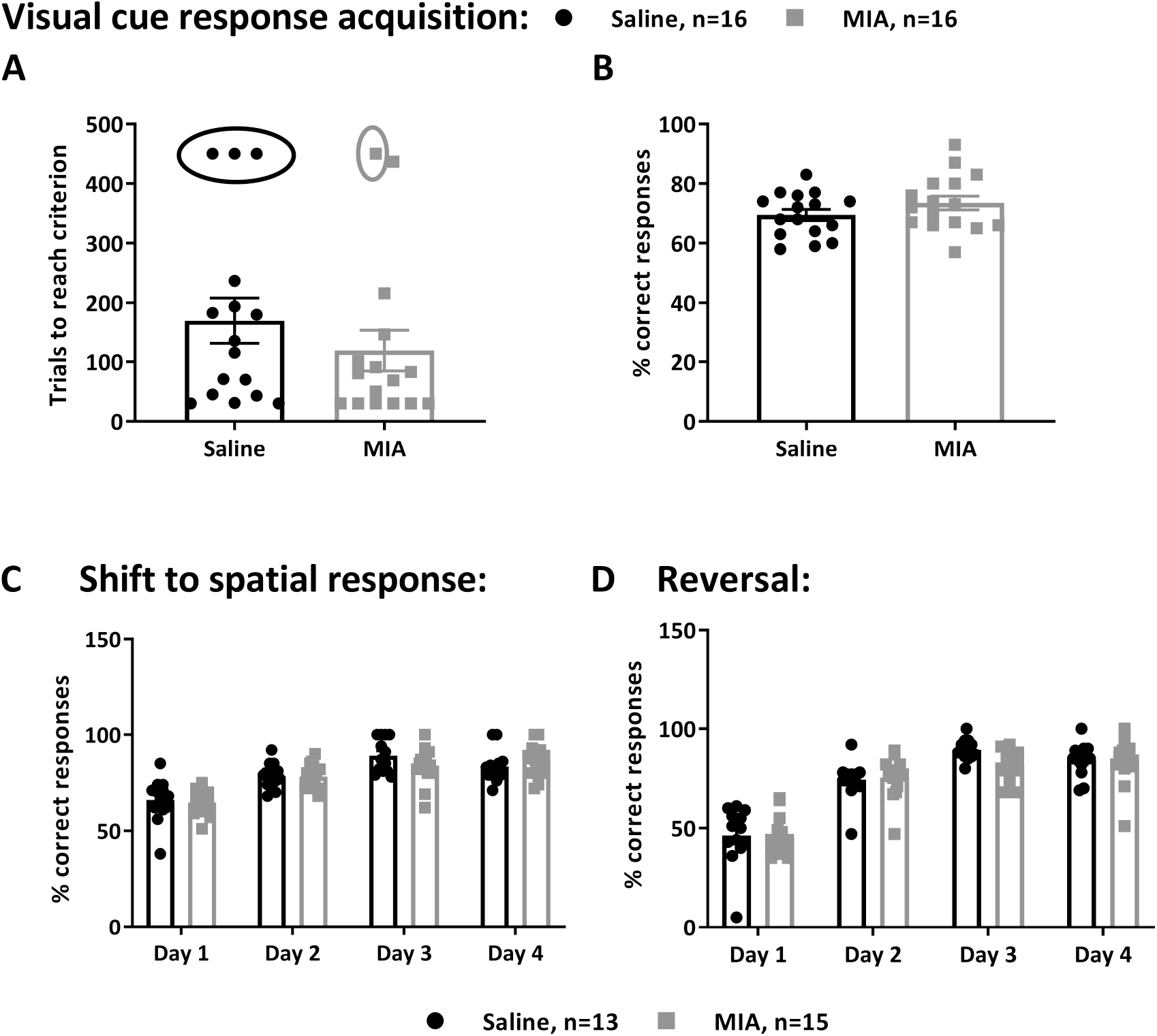
Intact performance on the operant task of behavioural flexibility after MIA model induction. Rats were injected with either 50ul of 3mg of MIA (▪, grey squares) or 50µl saline (•, black dots) in the le knee. Rats were tested on this task 38 days after model induction. Acquisition of the initial cue discrimination task, as reflected by **A**) trials to criterion and **B**) % of correct responses, was not affected by MIA injection compared with controls. Rats in the ellipses did not reach criterion on the 3 days of testing and, therefore, did not continue to the further testing phases. MIA- and saline-injected rats shied similarly well from the cue response rule to a spatial rule, as reflected by both groups showing similar **C**) % of correct responses across the 4 days of the shift phase. Similarly, both groups showed similarly efficient response reversals, as reflected by both groups showing similar **D**) % of correct responses. Data are presented as mean±SEM.

Shifting from visual cue discrimination to the spatial response rule was also not affected by MIA treatment. Both MIA- and saline-injected rats showed a similar number of trials to reach criterion on the first day (mean±SEM: saline, 111.9±12.4; MIA, 104.1+4.3; t_(26)_<1) and similarly increased their % of correct responses across the 4 days of the shift phase (main effect of day: F_(3,78)_=38.73, p<0.0001; no main effect or interaction involving group: F_(1,26)_<1 and F_(3,78)_>1.77, p>0.2, respectively; Figure 5C). Omissions were low (< 1 %) and did not differ between groups (data not shown).

Finally, MIA treatment did not impair the ability to reverse the spatial response rule. Both groups showed a similar number of trials to reach criterion on the first day (mean+SEM: saline, 111.2±6.3; MIA, 114.9+7.5; t_(26)_<1) and an increase in the % of correct responses across the 4 days of the reversal phase (time: F_(3,78)_=130.5, p<0.0001) (Figure 5D). However, for both measures, there was no main effect or interaction involving group (F_(3,78)_<1.67, p>0.2). Omissions were never greater than 1 omission per session and did not differ between groups (data not shown).

## 4. DISCUSSION

The MIA model was associated with OA-like knee pathology and knee pain, reflected by weight-bearing asymmetry, but no changes in PWTs, and reduced rearing behaviour in young adult male Lister hooded rats. MIA treated rats did not show any impairment in performance in memory tests (watermaze DMP test of rapid place learning; NOR test) dependent on the medial temporal lobe, including the hippocampus, or on an operant assay of behavioural flexibility, which has been associated with fronto-striatal mechanisms.

### MIA-induced knee pathology and pain phenotype in young adult LH rats

To our knowledge, this was the first study of the MIA model of OA pain in non-albino rats. We used Lister Hooded rats, because these are more suitable for cognitive assays than albino strains, which have been reported to perform poorly in watermaze and operant assays [20,29,66], probably partly reflecting their diminished vision [54]. MIA treatment produced consistent pain behaviour and knee pathology in Lister Hooded rats, including robust weight-bearing asymmetry and knee pathology with a loss of cartilage integrity, synovitis and occasional osteophyte formation. Following MIA model induction, rats also exhibited decreased rearing activity. Reduced rearing activity has previously been reported in MIA-injected Sprague Dawley rats and may reflect movement-provoked pain [42]. The slower swim speed in the MIA-treated rats compared with saline controls during watermaze testing at week 7 and 12 after MIA treatment may also reflect knee pain, although the reason for the swim speed increase in the control group is not clear. There were no consistent MIA-induced changes in horizontal locomotor activity, or in the acoustic startle response and its PPI.

In the present study, MIA treatment did not result in a change in PWT, contrasting with studies in Sprague Dawley rats [19,58]. Lowering of hindpaw withdrawal thresholds is often considered to indicate the presence of central sensitisation mechanisms, since it demonstrates enhanced mechanical sensitivity at a site distal to the injured joint [36]. Our data may, therefore, suggest that the MIA model of OA pain did not cause central sensitisation in Lister hooded rats. However, the sensitivity of the von Frey test to detect PWT reductions in Lister hooded rats is worth further consideration. Baseline PWTs were lower in Lister hooded rats (PWT log=0.8-0.9) than in SD rats (around PWT log=1.25 [32]), and, therefore this measure of pain behaviour is more sensitive to a floor effect in Lister hooded rats. In addition, Lister hooded rats are very inquisitive, which although beneficial for cognitive testing, makes von-Frey testing challenging. In the present study, although Lister hooded rats took longer to habituate to the testing environment, we were still able to measure PWTs. Another recent study recently reported difficulties with measuring PWTs in a model of neuropathic pain (partial saphenous nerve injury) in Lister hooded rats, but the authors were able to obtain measurements in the majority of rats and reported reduced PWTs [51]. Thus, on balance, it seems probable that the lack of effect of the MIA model on PWTs in Lister hooded rats reflects a lack of central sensitisation, rather than a consequence of a technical limitation.

### No effect of the MIA model on hippocampus-dependent memory

In the present study, neither rapid place learning performance in the watermaze DMP task nor NOR memory was altered by the MIA model. Previously, impaired watermaze place learning performance and NOR memory has been reported in rodent models of neuropathic and inflammatory pain [39,67]. Impaired NOR memory was also reported in MIA treated mice [43], contrasting with our finding of intact NOR memory in MIA treated rats. The difference between the MIA model of OA pain in Lister hooded rats and these other studies may be the level of pain intensity and/or the involvement of different brain regions, including hippocampal changes. Indeed, MIA-treated mice show markedly reduced PWTs [43], which likely reflects higher pain intensity and stronger involvement of central sensitisation in the mouse model. Although hippocampal changes in the MIA model have not yet been examined, studies have reported changes in the hippocampus of Sprague-Dawley rats with models of neuropathic and inflammatory pain [55,56,67]. It is noteworthy that a study conducted in Wistar Han rats (3-, 10- and 22-months old) with spared nerve injury did not report impairments in watermaze place learning performance [31], indicating that effects on this measure of hippocampal function are not always observed in neuropathic pain models.

Our findings of intact hippocampus-dependent memory in MIA-injected Lister hooded rats are consistent with the suggestion from human imaging studies that other pain conditions, including chronic back pain and complex regional pain syndrome, affect the hippocampus more than chronic pain associated with OA-related knee damage. More specifically, chronic back pain and complex regional pain syndrome, but not knee OA, were reported to be associated with reduced hippocampal volume [41]. In addition, relatively intact recognition memory in some chronic pain conditions, including chronic back pain and fibromyalgia, has been reported [30,59], supporting limited susceptibility of recognition memory to some chronic pain conditions.

### No effect of the MIA model on the operant task of behavioural flexibility

In the present study, MIA treatment did not impair operant performance and operant measures of behavioural flexibility, including rule shifting and reversal learning. Previous studies in rat models of neuropathic pain reported impaired operant performance, including slower adaptation to optimal choices [15], and impaired spatial reversal learning in the watermaze [31]. These measures do appear to have translational relevance as chronic pain patients with fibromyalgia and primary Sjögren syndrome with small fibre neuropathy show impaired behavioural flexibility on the Wisconsin Card Sorting Test [22,63]. In addition, in older adults with mixed pain conditions, pain severity (measured by questionnaire) was associated with impaired mental flexibility measured with the Trail Making Test [26]. The absence of impairments in operant performance and behavioural flexibility in MIA-injected Lister hooded rats, which contrasts with impairments in some chronic pain conditions reported by previous clinical and pre-clinical studies, may reflect lower severity of or more limited involvement of central mechanisms in the chronic pain associated with OA-like knee pathology in the present study compared to the other chronic pain conditions.

### Chronic OA pain and cognitive deficits: translational relevance

In the present study, we used an established model of OA pain in a rat strain well-suited for the investigation of cognitive function. In our study, OA-like pain behaviour was not associated with cognitive impairment in LH rats, unlike other experimental chronic pain conditions. The absence of impairments in hippocampal learning and memory function in the present study is consistent with human neuroimaging findings, suggesting that knee OA impacts the hippocampus less than other chronic pain conditions [41]. However, there is also evidence that chronic pain caused by knee OA causes forebrain changes [4,5,23], and recent studies in human participants support that OA pain may contribute to cognitive impairments, reflected in measures of verbal recall and fluency, subtraction ability and attention [24,27]. These measures may partly depend on similar neurocognitive mechanisms as those assessed in the present study, although it remains to be examined if human participants with chronic OA pain are impaired on translational behavioural assays that correspond more directly to the ones used in the present study, such as a virtual DMP test [10] and CANTAB assays of executive functions, including behavioural flexibility [3]. There are several factors that may interact with chronic pain to impair cognitive function in patients, and which impact upon the translational relevance of the experimental models. First, ageing is a key contributor to cognitive decline [16]. This is of relevance as OA is an age-associated disorder [13], and the average age of participants in the two recent studies showing a negative impact of chronic OA pain on cognition in human participants was > 60 years at the start of data collection [24,27]. Therefore, ageing may interact with chronic OA pain to impair cognitive function. In support of this possibility, impairments in behavioural flexibility associated with chronic pain have been reported to be age-dependent in both rodent [31] and human studies [64]. Moreover, recent neuroimaging evidence suggests that ageing may increase the vulnerability of a brain circuit including PFC and hippocampus to chronic lower back pain [2]. suggesting that the cognitive functions associated with these brain regions may also become more vulnerable to chronic pain with age. Second, social deprivation and education have recently been reported to interact with chronic pain to affect cognitive ability in people with OA [27]. Third, prescribed OA pain medication may contribute to cognitive impairments [40]. In particular, a negative impact of long-term opioid treatment on cognitive task performance has been reported in patients with chronic pain [57,59]. Examining the impact of age and pain medication on cognitive task performance in the MIA rat model offers the opportunity to test directly how these factors interact with chronic pain caused by OA-like knee damage to cause cognitive impairment, whereas the impact of socio-economic factors cannot be modelled experimentally in animals.

## ACKNOWLEDGMENTS

The authors thank the following people for their contributions to this article: Seyed Shahtaheri for helping with knee pathology scoring. This work was supported by Versus Arthritis United Kingdom (grant 20777). Author contributions: Study concept and design: S.G., G.J.H., S.G.W., V.C., and T.B.; Data acquisition: S.G.; Data analysis: S.G., G.J.H., V.C., and T.B; Manuscript-original: S.G., V.C. and T.B. Manuscript-review and editing: G.J.H. and S.G.W. All authors have seen and approved the final version of the article. Conflict of Interest: The authors declare no competing financial interests.

## Notes

### Competing Interest Statement

The authors have declared no competing interest.

