## Supplementary Material for "No evidence for cognitive impairment in an experimental rat model of knee osteoarthritis and associated chronic pain"

Contents:

1. Supplementary methods – detailed description of apparatus and procedures used for measurements of learning and memory and of behavioural flexibility

2. Supplementary figures and results

### 1. Supplementary methods – detailed description of apparatus and procedures used for measurements of learning and memory and of behavioural flexibility

#### 1.1. Watermaze delayed-matching-to-place (DMP) task

The watermaze DMP task is highly sensitive to hippocampal dysfunction [1,6,7,10]. The watermaze DMP task requires rats to learn rapidly, within one trial, the daily changing place of a hidden platform to escape efficiently from a circular pool of water, which is surrounded by prominent visual cues for spatial orientation. Rats were pre-trained on the watermaze DMP task, and their baseline performance was measured before MIA model induction. After model induction, watermaze DMP performance was evaluated at weeks 2, 4, 7 and 12 (Figure 1A). Methods were adapted from previous studies [1,6,7,9,10].

APPARATUS: The watermaze consisted of an open-field circular white pool (2 m diameter and 60 cm height) filled with water at 25±1°C that made opaque by the addition of children's white paint (Go Create, Tesco, UK). Four start points were evenly spaced along the circumference of the pool (north [N], east [E], south [S], and west [W]). Hidden in the watermaze pool (1–3 cm below the water surface) was an escape "Atlantis platform" (Med Associates Inc., US), which can be withheld at >20 cm below the water surface by a computer-controlled electromagnet for a predetermined time, making it unavailable to the rats. The room was brightly lit during watermaze testing, with a light level of about 200 lux at water level. The room was filled with spatial cues, including a traffic cone, lampshades hanging from the wall and different geometric 2D and 3D shapes, which were placed at different distances around the pool to aid spatial orientation. The rats' behaviour was recorded by an overhead video camera connected to a computer with a programme for video capture and with EthoVision XT 8.5 software (Noldus, Netherlands) in an adjacent control room. The EthoVision software can compute various behavioral measures, including latencies and path lengths to reach the platform location, and times in different areas of the pool.

PROCEDURE: Rats performed four daily trials. During trial 1, rats can rapidly learn the novel location of the hidden platform, and then on subsequent trials they can use the place memory to efficiently locate the hidden platform. To start the test, rats were placed into the water facing the pool walls at one of the four start positions. A different start position was used for each of the four daily trials, in a predetermined and arbitrary sequence, to prevent the use of egocentric strategies. The platform location was changed daily, but remained constant during the four consecutive daily trials. Rats were tested with a novel platform location each day. The centre of the escape platform was either located on an inner (0.8 m) or outer (1.4 m) ring concentric with the pool. Each trial had a maximum duration of 120 s, after which the rat was guided to the platform by the experimenter, if it had not found the platform by itself. Following each trial, rats were allowed 30 s on the platform, before they were dried gently on a towel and placed in a holding box while the experimenter was setting up the next trial, resulting in an inter-trial interval of around 10-30 s. Trial 2 was occasionally run as “probe trial”. During this probe, the platform was withheld for the first 60 s, to monitor the rats’ search preference for the zone containing the platform (the target zone). After the 60 s, the platform was automatically released allowing the rat to find and climb onto it. Between trial 1 and the probe trial the inter-trial interval was 20 min, instead of 10-30 s, because this renders the task more sensitive to any impairment in hippocampal plasticity mechanisms [10].

Before model induction, rats were pre-trained on the watermaze DMP task and baseline performance was assessed (Figure 1A). Pre-training consisted of 8 consecutive days of DMP testing, with trial 2 on days 6 and 8 run as probe trials. The retention delay between trials 1 and 2 was 10-30 s on the first four days of pre-training and 20 min on the probe days and the preceding standard training days. After MIA model induction, watermaze DMP performance was re-assessed at weeks 2, 4, 7 and 12 (Figure 1A). In each week, rats underwent 4-day testing blocks, consisting of two probe days (second and fourth day) to obtain search preference measures of 1-trial place learning, with each probe day preceded by one day of testing with 4 standard trials. The retention delay between trial 1 and 2 during these 4-day test blocks was 10-30s on standard testing days and 20 min on probe days.

PERFORMANCE MEASURES: Search preference for the platform area and its vicinity was used as main performance measure of rapid, 1-trial, place memory performance. Based on previous studies, search preference is more dependent on the hippocampus than latency and path length measures [1,6,7], which are also associated with higher variability [9] (also see [2] for related findings in human participants). To quantify search preference for the correct zone, eight virtual 20 cm diameter zones were defined on the inner and outer ring of the pool, so that one zone, the “correct” zone was concentric with the platform location, and all 8 zones were non-overlapping. The time spent in each of these 8 zones during the 60 s probe trial was determined automatically using the Ethovision software. Percentage of time searching the correct zone was calculated as: [time in correct zone (s) / total time in all 8 zones (s)] x 100. By chance, this value should be 100%/ 8 = 12.5%, whereas higher values indicate a search preference for the correct zone. Additionally, latencies and path lengths to reach the platform perimeter were recorded for all trials, with steep reduction from trial 1 to trial 2 indicating 1-trial place memory. Path lengths were analysed instead of latencies following MIA model induction, because they have the advantage over latencies that they measure the efficiency in reaching the platform independent of potential model-induced swim speed changes. Swim speed was also measured during the 60-s probe trials, to examine if MIA-injected rats developed any motor impairments.

For analysis, search preference and swim speed was averaged across the two probes at each testing time point (on day 6 and 8 for baseline testing and the two probes conducted as part of the 4-day testing blocks at each post-operative testing time point) and path length measures for trial 1 to 4 were averaged across all four days at each testing time point (days 4 to 8 of pretraining for baseline values and all four days of the 4-day test blocks at all post-operative testing time points).

### 1.2. Novel object recognition (NOR) memory test

The NOR memory test exploits rats' innate tendency to explore novel stimuli, including novel objects, more than familiar stimuli [5,8]. During the test, rats are presented with a novel object and a familiar object, i.e. an object that they had the opportunity to explore during a preceding sample phase. Rats will typically spend more time exploring the novel object, which reflects recognition memory of the familiar object that the rats explored during the sample phase. The NOR test allows to evaluate object recognition memory without use of training or reinforcement. Brain areas required for simple NOR memory include medial temporal lobe regions, especially the perirhinal cortex, whose key role in NOR memory has been long established [12], and also the hippocampus, especially if the retention delay between sample phase and test is greater than 10 min [4]. In study 2, we assessed object recognition memory at baseline, before MIA model induction, and on day 29-31 after model induction (Fig. 1B), using an NOR memory test procedure with a 24-h retention delay between sample phase and test, which was adapted from [8].

APPARATUS: Rats were tested in groups of 4 simultaneously in individual plastic rectangular arenas (38 x 40 x 54 cm high walls) with an opaque plastic lid, which were placed in a well-illuminated room (200 lux). Objects consisted of duplicate copies of glass or plastic bottles with different shapes, colours and sizes. These objects were filled with water to make them too heavy to be displaced by the rat. Objects were counterbalanced across groups and placements (right or left of arena). Sessions were recorded using an overhead camera and later analysed. Arenas and objects were cleaned with 20% ethanol before each trial/session to remove any odour cues the rats may have left.

PROCEDURE: NOR testing consisted of habituation, sample and test phases, performed across 3 consecutive days. Habituation: On the first day, rats were placed into the empty arena for 1h of habituation, to familiarize them with the test arena. Sample phase: On the following day, rats were placed in the empty arenas for 3 min of re-acclimatisation, returned to the home cage for 30-45 s while arenas were cleaned and objects put in their positions, and then re-placed in the arenas for the sample phase, during which they could 'familiarise' themselves with one object. The two copies of this to be 'familiar' object were placed in opposite corners of the arena and rats were then allowed to explore them for 5 min, after which they were returned to the home cage. Test phase: After a retention delay of 24 h, rats were replaced into the arena for 3 min; at this point, the arena contained one copy of the familiar object used in the sample phase and one "novel" object. Both sample and testing phases were recorded and later analysed.

PERFORMANCE MEASURES: Time exploring each object was defined as only direct contact with or active exploration of the object by directing the nose towards the object from a distance of less than 1 cm, e.g. sniffing and or interacting with the object. Contact with the object, but not facing it from a distance of more than 1 cm or sitting next to it, was not scored as exploration time. The discrimination ratio D was calculated as follows: D = total time exploring novel object/ (total time exploring novel object + total time exploring familiar object) [5,8].

#### 1.3. Operant test of behavioural flexibility

An operant lever press task adapted from [3], was used to assess behavioural flexibility - the ability to switch stimulus-response patterns rapidly, as reflected by response shifts or reversals [11]. Shifts refer to a subject beginning to make responses to new or previously irrelevant stimuli (e.g., from pressing the lever indicated by a cue light to pressing either the left or right lever in an operant box), whereas reversals refer to changing a response from a previously rewarded to a previously non-rewarded stimulus within the same category (e.g., from left to right lever). Previous studies suggest that behavioural flexibility as measured on the operant task depends on the PFC and subcortical regions, including ventral striatum [3]. In study 2, following MIA model induction, rats were trained first to acquire a lever press response to a cue light to receive food reward, before being tested for their ability to shift this response to a spatial response (pressing either the left or right lever to receive food) and then to reverse this spatial response (start to press the opposite lever to received food reward) (Fig. 1B). Operant testing procedures were adapted from previous studies [3].

APPARATUS: The task was conducted using eight individual operant chambers (Med Associates Inc., US). Each chamber was equipped with a house light, two retractable levers, two stimulus lights above the levers and a reinforcement pellet dispenser located between the levers. Each rat was assigned to an operant chamber, where it underwent all operant test sessions. Chambers were cleaned with 20% ethanol between different rats. The stimuli presented, lever operation and data collection were controlled via an interface with the computer and using custom software (MED-PC software) [3].

PROCEDURE: Before testing on the food-reinforced operant task, rats were put on a restricted diet to motivate responding. Food-restriction was gradually introduced a few days prior to the beginning of the pretraining. The target weight was 85-90% of the free feeding weight, based on a pre-established weight growth curve. Rats were weighed every day before the task and received their normal food in their home cages; on test days, rats received their food after completing the day's operant task session.

Rats were first pretrained across 3 phases to familiarise them with the task apparatus and with pressing both levers; on the day before pretraining, 10-20 sugar pellets (Purified rodent tablet 5TUL, TestDiet, US) per rat were placed in the rats' home cage to familiarise them with the pellets. Phase 1: In this phase, the food restricted rats were trained to press an extended lever, and one reward pellet was delivered for each lever press. Rats completed one 30-min session per day for four days. Phase 1 had a duration of 4 days. On the first 2 days, rats were trained on one of the two levers (right or left) and on the next 2 days they were trained on the opposite lever, with half of the rats first trained on the right lever and the other half on the left lever. On the first day only, two reward pellets were placed in the magazine cup and crushed pellets on the top of the extended lever at the beginning of the session, so rats readily approached these components of the test box. At the end of this phase, all rats made a minimum of 50 lever presses per session. Phase 2: Rats were then trained on the retractable lever to familiarise them with the extension and retraction of the levers and the associated sound. Levers were pseudorandomly extended, but the same lever was not presented more than two consecutive times. Rats were placed into chambers with the house light off. As soon as the experimenter started the program, both stimulus lights turned on, and 3 s later the house light came on and one of the levers extended for 10 s. Pressing the lever resulted in its retraction, release of a reward pellet and switching off of all lights. If the rat did not press the lever, this was considered an omission. Each session consisted of 90 trials. Phase 2 lasted 5 days, and on the 5th day all rats made fewer than 5 omissions over the session. Phase 3: This started immediately after the last session of phase 2 was completed. The rats remained in the chamber and were assessed for their side preference. Phase 3 consisted of seven trials, each of which was composed of between two and eight sub-trials (20 s sub-trial interval). On each sub-trial, both levers were extended into the chamber for 10 s or until a lever press response, while the stimulus lights were off. A response on either lever on the first sub-trial of each trial was rewarded, and recorded as the "initial response". A response on the same lever on the subsequent sub-trials within the same trial was not rewarded. Six subsequent responses on the same lever with a trial were allowed, before the rats were given a forced-choice sub-trial with only the opposite lever extended. After the initial response, the opposite levers should be pressed 7 times, so 7 + 7 presses in total. Phase 3 ended after rats had completed 7 pairs of rewarded trials.

Following pretraining, rats underwent three testing phases, including visual cue discrimination learning, a rule shift to a spatial response (left or right lever press) task, and, finally, a phase involving reversal of the spatial response. Phase 4 (Visual cue discrimination learning): Both levers were extended into the chamber and only one cue light was illuminated. To receive a reward, the rat had to press the lever with the cue light illuminated above it. An individual session stopped after the rat reached the criterion of 10 consecutive correct trials or after a maximum of 150 trials. If rats did not reach the criterion on day 1, the rat was tested again on the following day, up to a maximum of 3 days in total. Rats that did not reach the criterion on the 3rd day were excluded from further behavioural testing.

Phase 5 (Shift from visual cue to spatial response): Phase 5 began with 20 trials that were identical to those of phase 4, i.e. responding according to the cue rule was rewarded. These trials served to measure retrieval/expression of the cue rule – no differences were detected. On trial 21, the program shifted to the spatial response rule. The left or right stimulus light was pseudorandomly illuminated for 3 s, then both levers extended into the chamber for 10 s or until a response occurred. The reward was delivered only when the rat pressed the opposite lever of the side bias defined during side preference training (phase3), independent of the position of the cue light. The session stopped after the rat reached the criterion of 10 consecutive correct responses and only after it had completed a minimum of 30 trials, or after a maximum of 180 trials (reminder trials included) on the first set shift day and a maximum of 150 trials on subsequent set shift days. Rats performed 3 consecutive days of this task. A few days later, a fourth day as a reminder of phase 5 was presented to the rats, to ensure robust performance on the spatial response task, before rats underwent testing that required reversals of the spatial response.

Phase 6 (Spatial response reversal): The reversal task was run similar to phase 5. A session started with the left or right stimulus light being pseudorandomly illuminated for 3 s, then both levers were extended into the chamber for 10 s or until a response occurred. However, here the correct response was ‘reversed’ compared to phase 5. So, if in phase 5 the designated lever for a particular rat was the right lever, now the left lever was the designated lever to receive a reward pellet, and vice versa. The criterion was again 10 consecutive correct responses, with a maximum of 150 trials. All rats performed this task for 4 consecutive days, with the same lever remaining the correct lever throughout the 4 days.

PERFORMANCE MEASURES: The analysis of this test focused on trials to criterion, the percentage of correct responses and percentage of omissions.

### Supplementary figures and results

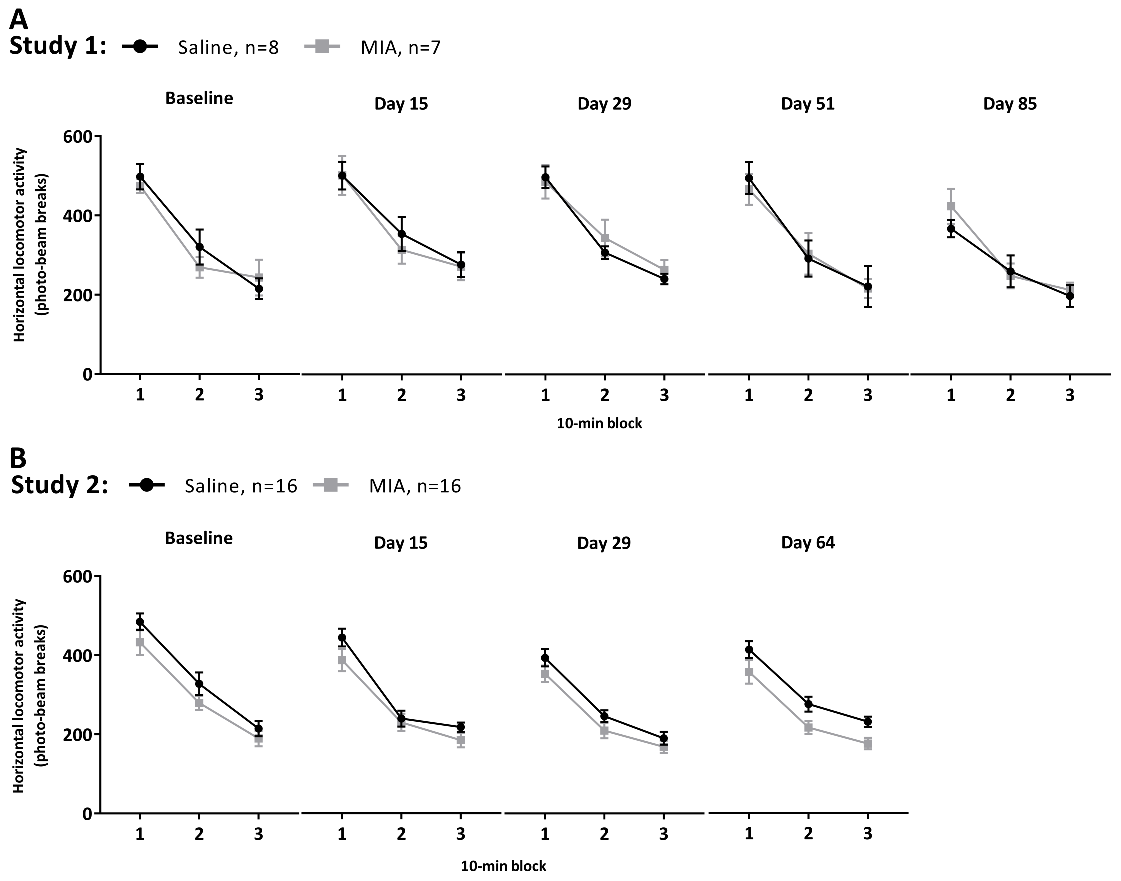


**Figure S1 – MIA injection in Lister hooded rats did not affect horizontal locomotor activity.** Rats were injected with either 3mg/50µl MIA (▪, grey squares) or 50µl saline (●, black dots) in the left knee. **A)** study 1 - group: $F_{(1,13)}$=0.02, p=0.88; time: $F_{(4,52)}$=6.05; p<0.0001; block: $F_{(2,26)}$=146.45; p<0.0001; block x group: $F_{(2,26)}$=0.16, p=0.85; time x group: $F_{(4,52)}$=0.85; p=0.50; block x time: $F_{(8,104)}$=1.16; p=0.39; block x time x group: $F_{(8,104)}$=0.43; p=0.90. **B)** study 2 - group: $F_{(1,30)}$=3.65, p=0.07; time: $F_{(2,60)}$=3.42; p=0.04; Block: $F_{(2,60)}$=319.19; p=0.0001; time x group: $F_{(2,60)}$=0.60; p=0.56; block x group : $F_{(2,60)}$=1.05; p=0.36; time x block x group: $F_{(4,120)}$=0.92; p=0.05. Mixed factorial ANOVA. Data are presented as mean±SEM.

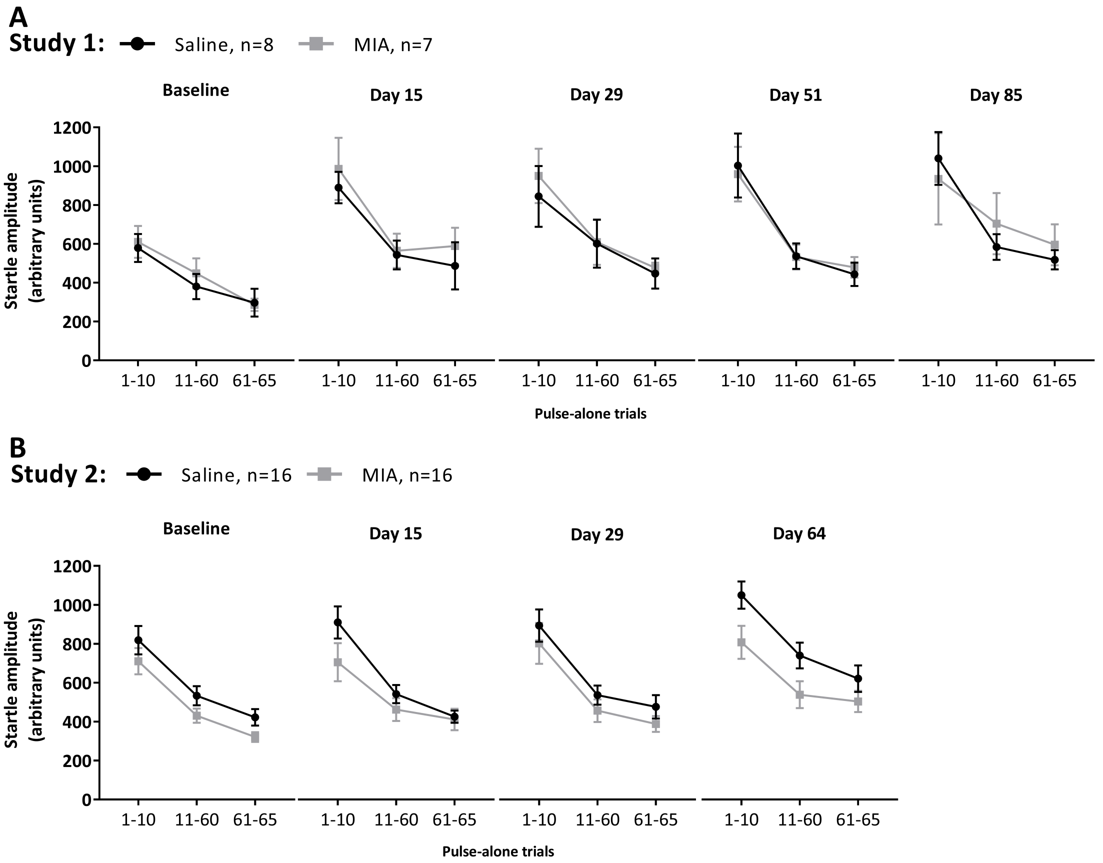


**Figure S2 – Startle response was not affected by MIA-model induction.** Rats were injected with either 50ul of 3mg of MIA (▪, grey squares) or saline (●, black dots) in the left knee. **A)** study 1 - group: $F_{(1,13)}$=0.168; p=0.69; time: $F_{(4, 52)}$=8.028; p<0.0001; block: $F_{(2, 26)}$=70.453; p<0.0001; block x group: $F_{(2, 26)}$=0.056; p=0.945; time x group: $F_{(4, 52)}$=0.169; p=0.953; block x time: $F_{(8, 104)}$=1.605; p=0.132; block x time x group: $F_{(8, 104)}$=0.602; p=0.774. **B)** study 2 - group: $F_{(1,30)}$=2.91; p=0.1; time: $F_{(2,60)}$=8.99; p=0.000; block: $F_{(2,60)}$=113.06; p=0.000; time x group: $F_{(2,60)}$=1.26; p=0.29; block x group: $F_{(2,60)}$=1.88; p=0.16; block x time: $F_{(4,120)}$=0.37; p=0.83; time x block x group: $F_{(4,120)}$=0.78; p=0. 35. Mixed factorial ANOVA. Data are presented as mean±SEM.

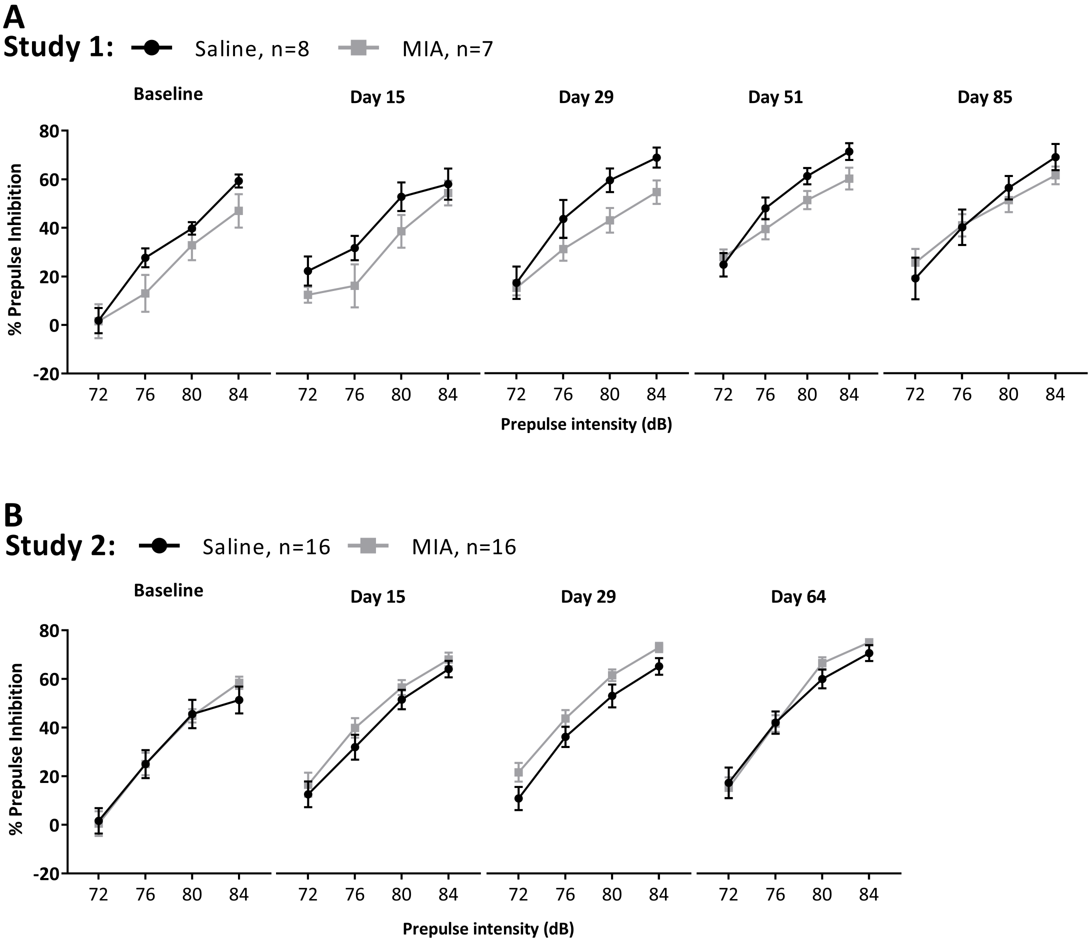


**Figure S3 – Prepulse inhibition of the startle response was not affected by MIA-model induction.** Rats were injected with either 50ul of 3mg of MIA (▪, grey squares) or saline (●, black dots) in the left knee. **A)** study 1 - group: F(1,13)=2.079; p=0.17, prepulse x time: F(12,156)=2.056; p=0.02, time x group: F(4,52)=0.82; p=0.52). **B)** study 2 - group: $F_{(1,30)}$=2.91; p=0.10; time: $F_{(2,60)}$=3.23; p=0.046; pulse: $F_{(3,90)}$=320.68; p=0.0001; time x group: $F_{(2,60)}$=0.98; p=0.38; pulse x group: $F_{(3,90)}$=0.15; p=0.93; pulse x time: $F_{(6,180)}$=0.11; p=0.39; time x pulse x group: $F_{(6,180)}$=0.87; p=0. 52. Mixed factorial ANOVA. Data are presented as mean±SEM.

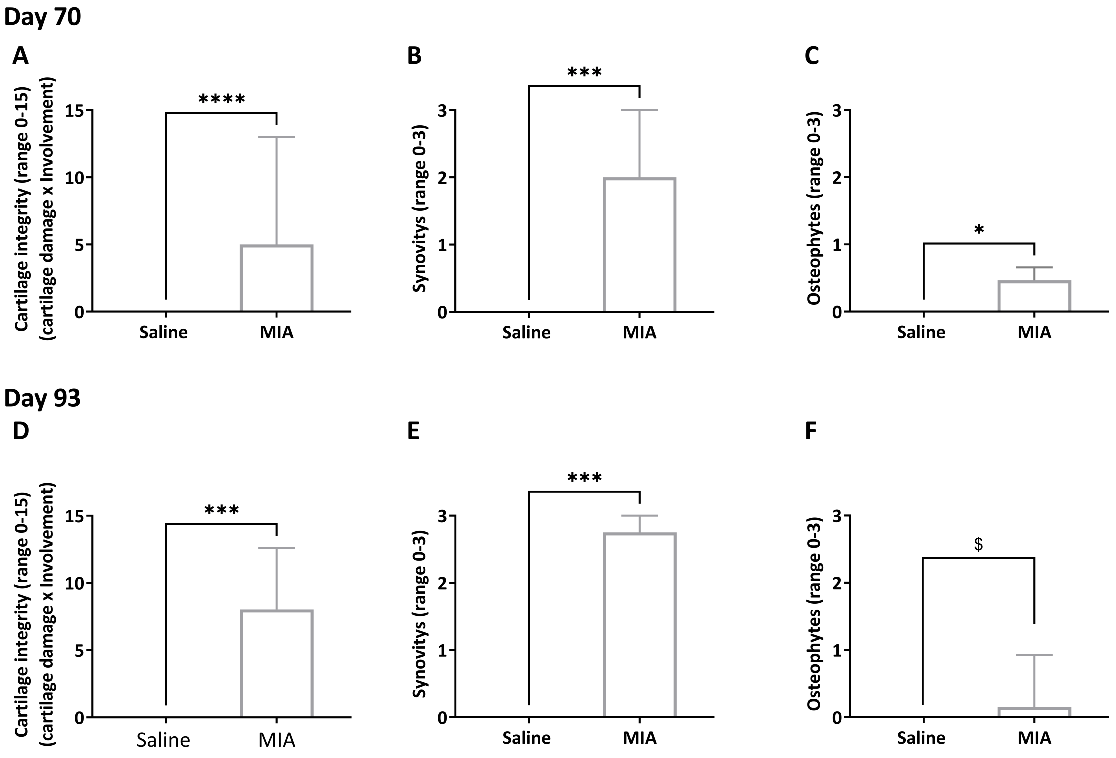


**Figure S4 – MIA-induced OA-like knee pathology in Lister hooded rats: Microscopic quantification of histological changes of tibial plateau (Study 1 and 2).** Rats were injected with either 50ul of 3mg of MIA (▪, grey) or saline (●, black) in the left knee. Knees were collected at the end of study 1 on day 93 after injection (MIA=7, saline=8) and study 2 on day 70 after injection (MIA=16, saline=16), and knee pathology was examined. There was similarly pronounced OA-like knee pathology in both studies. In knees collected on day 70 after model induction (study 2), MIA-injected rats showed marked **A**) cartilage damage, **B)** synovial inflammation, and a trend for higher number of **C)** osteophytes compared to saline controls. In knees collected on day 93 after model induction (study 1), MIA-injected rats showed **D**) cartilage damage, **E**) synovitis and **F)** osteophyte formation. Data are presented as mean±SEM, where assumptions of normality were not violated (A and F) or as median±interquartile range, where assumption of normality were violated (B, C, D and E). *p<0.05, **p<0.001, ***p<0.0001, $p<0.1, 2-way ANOVA with Bonferroni multiple comparisons post-hoc testing.
